# The inhibition of cellular toxicity of amyloid-beta by dissociated transthyretin

**DOI:** 10.1101/852715

**Authors:** Qin Cao, Daniel H. Anderson, Wilson Liang, Joshua Chou, Lorena Saelices

## Abstract

The protective effect of transthyretin (TTR) on cellular toxicity of amyloid-beta (Aβ) has been previously reported. TTR is a tetrameric carrier of thyroxine in blood and cerebrospinal fluid, whose pathogenic aggregation causes systemic amyloidosis. In contrast, many reports have shown that TTR binds amyloid-beta (Aβ), associated with Alzheimer’s disease, alters its aggregation, and inhibits its toxicity both *in vitro* and *in vivo*. In this study, we question whether TTR amyloidogenic ability and its anti-amyloid inhibitory effect are associated. Our results indicate that the dissociation of the TTR tetramer, required for its amyloid pathogenesis, is also necessary to prevent cellular toxicity from Aβ oligomers. These findings suggest that the Aβ binding site of TTR may be hidden in its tetrameric form. Aided by computational docking and peptide screening, we identified a TTR segment that is capable of altering Aβ aggregation and toxicity, mimicking TTR cellular protection. This segment inhibits Aβ oligomer formation and also promotes the formation of non-toxic, non-amyloid, amorphous aggregates which are more sensitive to protease digestion. This segment also inhibits seeding of Aβ catalyzed by Aβ fibrils extracted from the brain of an Alzheimer’s patient. Our results suggest that mimicking the inhibitory effect of TTR with peptide-based therapeutics represents an additional avenue to explore for the treatment of Alzheimer’s disease.

**Significance statement:** The pathological landmarks of Alzheimer’s disease are the formation of amyloid plaques and neurofibrillary tangles. Amyloid plaques contain fibrous structures made of aggregated amyloid-beta (Aβ). In 1982, Shirahama and colleagues observed the presence of transthyretin (TTR) in these plaques. TTR is a tetrameric protein whose aggregation causes transthyretin amyloidosis. However, TTR protects Aβ from aggregating and causing toxicity to neurons. In this study, we show that the dissociation of TTR tetramers is required to inhibit cellular toxicity caused by Aβ. In addition, we identified a minimum segment of TTR that inhibits Aβ aggregation and cellular toxicity by the formation of amorphous aggregates that are sensitive to proteases, similar to the natural effect of TTR found by others in vivo.

## Introduction

The physiological importance of transthyretin in Alzheimer’s disease was first reported by Schwarzman et al. in 1994 (1). One of the hallmarks of Alzheimer’s disease (AD) is the formation of brain plaques composed of amyloid-beta peptide (Aβ). Forty-two-residue long amyloid-β peptide (Aβ42) is the predominant variant in neuritic plaques of AD patients, with higher amyloidogenicity and cellular toxicity *in vitro* (2, 3). Many studies have shown that transthyretin (TTR) binds to Aβ, alters its aggregation, and inhibits its toxicity both *in vitro* and *in vivo* (4–8). *In vitro*, TTR co-aggregates with Aβ oligomers into large non-toxic assemblies, thereby inhibiting cellular toxicity (9, 10). *In vivo*, TTR sequesters Aβ and facilitates its clearance in the brain (1). More recently, Buxbaum laboratory showed that overexpression of wild-type human TTR suppressed disease progression in the APP23 transgenic AD mouse model (6). They also showed that silencing the endogenous TTR gene in AD transgenic mice accelerated Aβ42 deposition (5).

What makes the pair of TTR and Aβ42 particularly interesting is the amyloid nature of the two elements. In health, TTR functions as a transporter of retinol and thyroxine in blood, cerebrospinal fluid, and the eye, and is secreted by the liver, choroid plexus, and retinal epithelium, respectively. However, dissociation of tetrameric TTR leads to amyloid fibril formation and systemic TTR amyloid deposition in patients of transthyretin amyloidosis (11–13). Whether the amyloidogenicity of TTR is linked to its interaction to Aβ42 is still under debate. Previous work showed that different aggregation propensities of TTR result in distinct interaction capabilities: less amyloidogenic variants had increased affinity for Aβ (7). However, this did not impact on the levels of inhibition of Aβ aggregation, and cytotoxicity protection was not assessed. Here we attempt to fill the experimental gap by studying the protective effect of TTR variants at different aggregated states over Aβ42 cellular toxicity. In addition, we show the identification and characterization of a segment of TTR that binds Aβ42 triggering the formation of non-amyloid amorphous aggregates that are more sensitive to protease digestion, therefore mimicking TTR inhibitory effect over Aβ42.

## Results

We first evaluated the protection against Aβ42 cytotoxicity by nine TTR variants with distinct aggregation propensities, in several aggregated states (Table 1). The amyloidogenic behavior of three representative TTR variants is shown in Fig 1A and 1B. For this assay, recombinant transthyretin was incubated at 37 °C, pH 4.3, protein aggregation was followed by immuno-dot blot of the insoluble fractions collected by centrifugation at several time points (Fig. 1A), and electron microscopy was performed after 4 days of incubation (Fig. 1B). We found that M-TTR, a monomeric-engineered form of transthyretin that is soluble at physiological pH (14), shows notable aggregation after one day of incubation at pH 4.3 (Fig. 1A-C). NSTTR is an artificially mutated variant of TTR carrying the double mutation N99R/S100R that, although remains tetrameric in solution (Fig. 1C), shows a slower aggregation pattern than M-TTR (Fig. 1A and 1B). T119M is a very stable mutant that did not show any sign of aggregation in four days (Fig. 1A-C). Consistently, T119M results in a significant delay of the onset of hereditary neuropathic ATTR in patients who carry both *ttr-V30M* and *ttr-T119M* genes (15). We then incubated Aβ42 with the samples obtained from Fig. 1A for 16 hours and evaluated its cytotoxicity by following MTT reduction. Three observations were made. The first observation is that once aggregated, TTR does not prevent Aβ42 toxicity, for any of the variants (Fig. 1D). In addition, the variant NS-TTR, which aggregates at a slow pace, resulted to be cytoprotective longer than M-TTR (Fig. 1D). The third observation is that T119M, which does not dissociate and aggregate, does not prevent toxicity either (Fig. 1D). We observed the same phenomena when other six TTR variants were evaluated (Supplemental Fig. 1). Although variants S112I, M13R/L17R, or A108R/L110R are mainly monomeric or dimeric in solution, the cytotoxic protection at day 0 is not greater than NSTTR or L55P that are mainly tetrameric in solution (Fig. 1C and Supplementary Fig. 1B). This finding suggests that the capacity to protect from Aβ42 toxicity does not correlate with the initial oligomeric state of soluble variants (Fig. 1C, and Supplemental Fig. 1B), but rather with the dissociation state at the moment of the co-incubation with Aβ42 (Fig. 1A and 1B, and Supplemental Fig. 1C). A control experiment shows that the TTR variants used in this assay were not cytotoxic in the absence of Aβ42, with the exception of M-TTR, which resulted in a 20% reduction of cell viability (Supplementary Fig. 1D). Overall, our results indicate that dissociation of TTR is required for the inhibition of Aβ cytotoxicity. In addition, the data suggest that a more stable dissociated TTR exerts more Aβ42 inhibition than a more amyloidogenic variant. These findings led us to wonder whether a segment of TTR that is only exposed when dissociated but not in the aggregate might be similarly capable of altering Aβ toxicity.

**Table 1.**
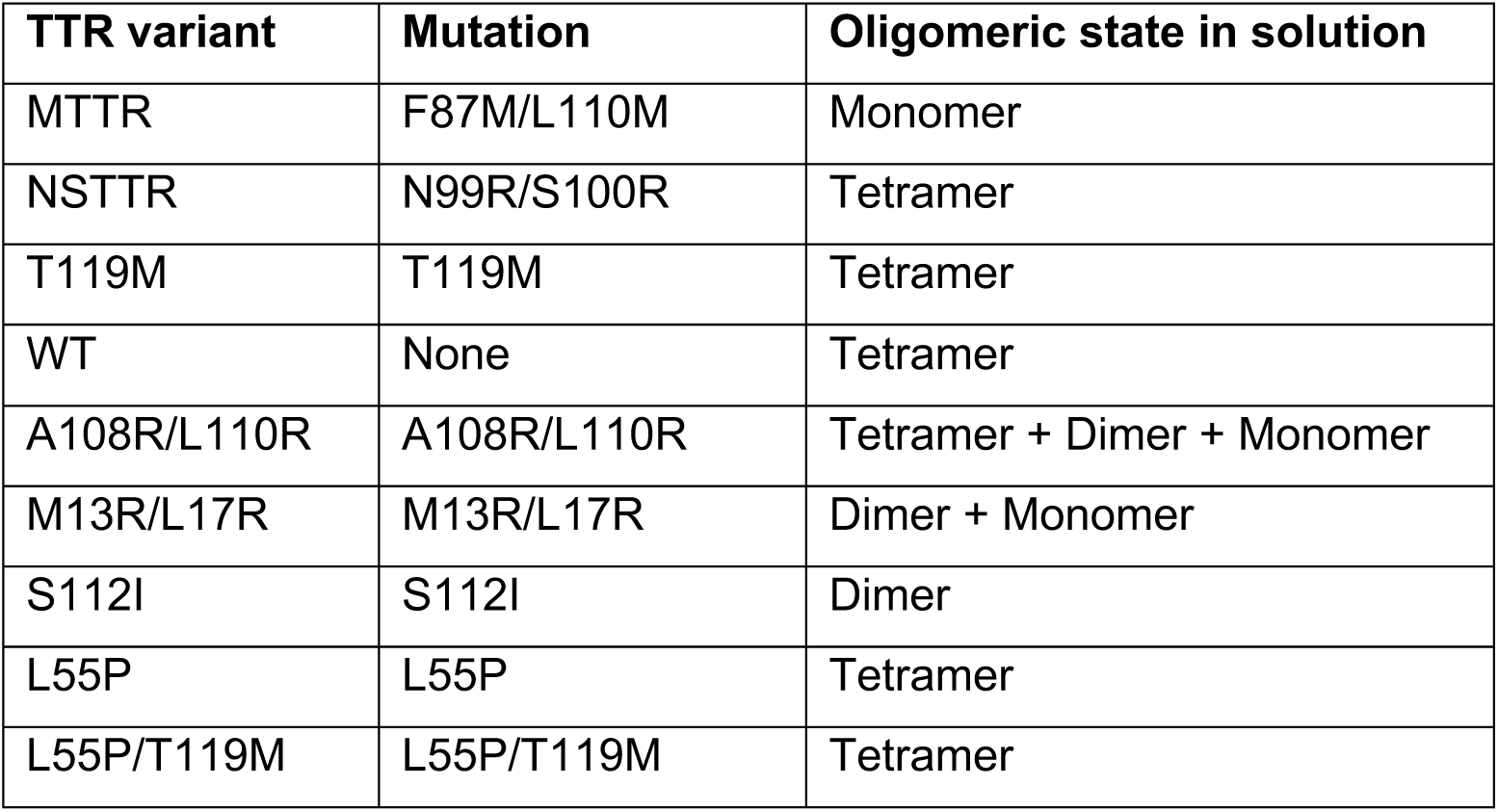
List of TTR variants evaluated in this study.

| **TTR variant** | **Mutation** | **Oligomeric state in solution** |
| --- | --- | --- |
| MTTR | F87M/L110M | Monomer |
| NSTTR | N99R/S100R | Tetramer |
| T119M | T119M | Tetramer |
| WT | None | Tetramer |
| A108R/L110R | A108R/L110R | Tetramer + Dimer + Monomer |
| M13R/L17R | M13R/L17R | Dimer + Monomer |
| S112I | S112I | Dimer |
| L55P | L55P | Tetramer |
| L55P/T119M | L55P/T119M | Tetramer |

**Figure 1.**
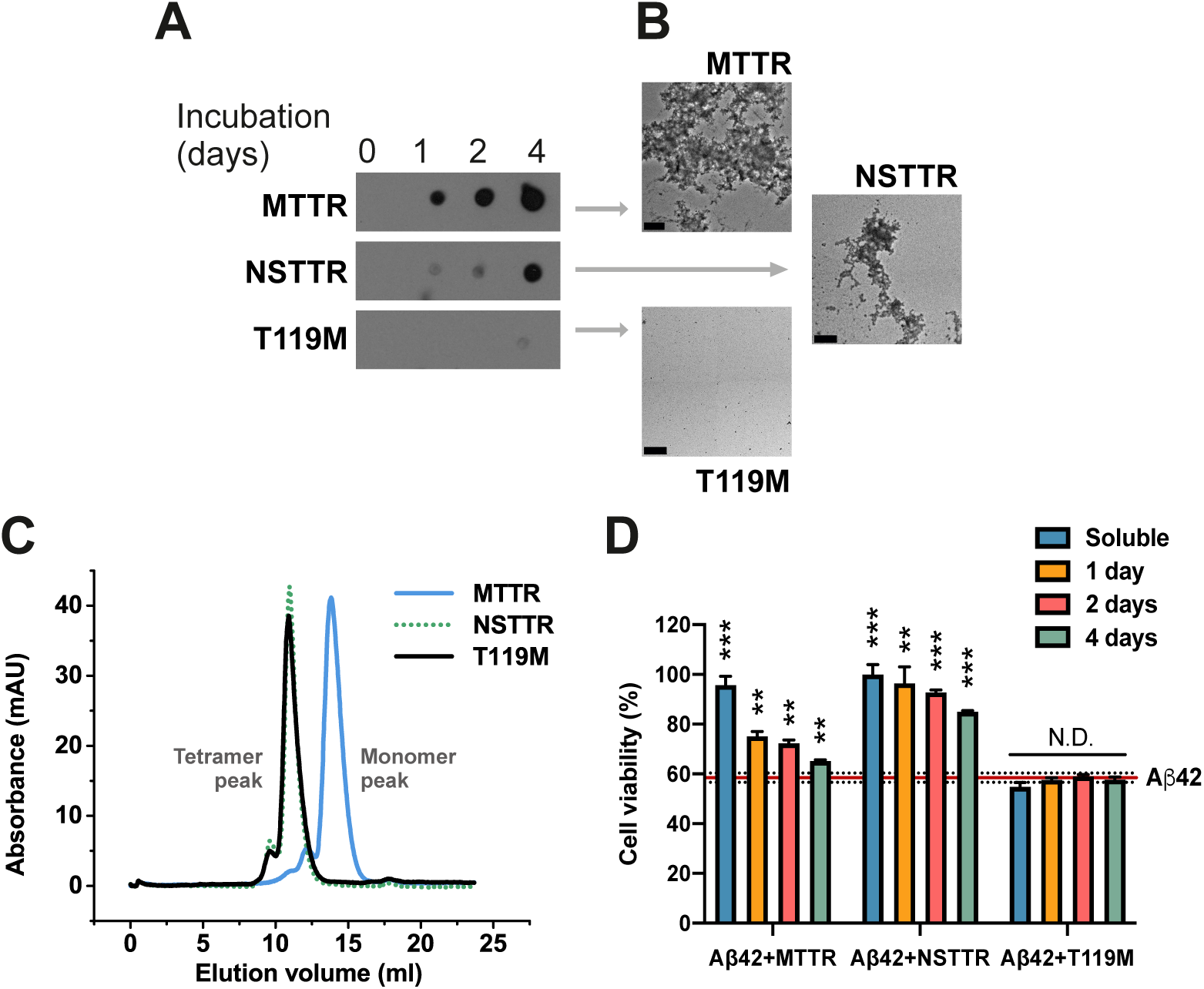
The inhibition of Aβ42 cytotoxicity by TTR depends on the aggregation propensity of each TTR variant. **(A)** Aggregation assay of three TTR variants followed by immunodot blot of insoluble fractions collected prior incubation, or after 1, 2, and 4 days of incubation at 37° C, as labeled. **(B)** Electron micrographs of aggregated TTR variants after 4 days of incubation. Scale bar, 200 nm. **(C)** Size exclusion chromatography of soluble TTR variants at pH 7.4. **(D)** Cytotoxicity assay of Aβ42 in the presence of TTR variants at different stages of aggregation, followed by MTT reduction. A five-fold molar excess of soluble TTR, and aggregated samples collected after 1, 2 and 4 days of incubation was added to soluble 10 μM Aβ42 and incubated overnight. Samples were added to HeLa cells, and MTT reduction was measured after 24 hours. Buffer-treated cells were considered 100% viability and used for normalization. n=3. Error bars, S.D. **p ≤ 0.005, ***p ≤ 0.0005, N.D., significance not detected. These results suggest that the dissociation of the tetrameric TTR structure precedes Aβ42 cytotoxicity protection, whereas aggregated TTR does not exert any effect.

Computational modeling of the interaction of a fibrillar segment of Aβ42 and the TTR monomer shows a tight packing of the thyroxine binding pocket and the fibrillar structure (Fig. 2). Others have shown that the amyloidogenic segment KLVFFA is protected when TTR is bound to Aβ (16). In previous studies, our lab determined the structures of three fibrillar polymorphs of KLVFFA by x-ray microcrystallography (17). We used these three polymorphs as well as the monomeric form of wild-type TTR (4TLT.pdb, (18)) to perform protein-protein computational docking (Fig. 2). For every fibrillar form, the segment TTR(105-117) was the longest interacting TTR segment. The segment TTR(105-117) is only exposed in the monomeric or dimeric form of TTR. We wondered if this segment might mimic TTR protective effects when isolated.

**Figure 2.**
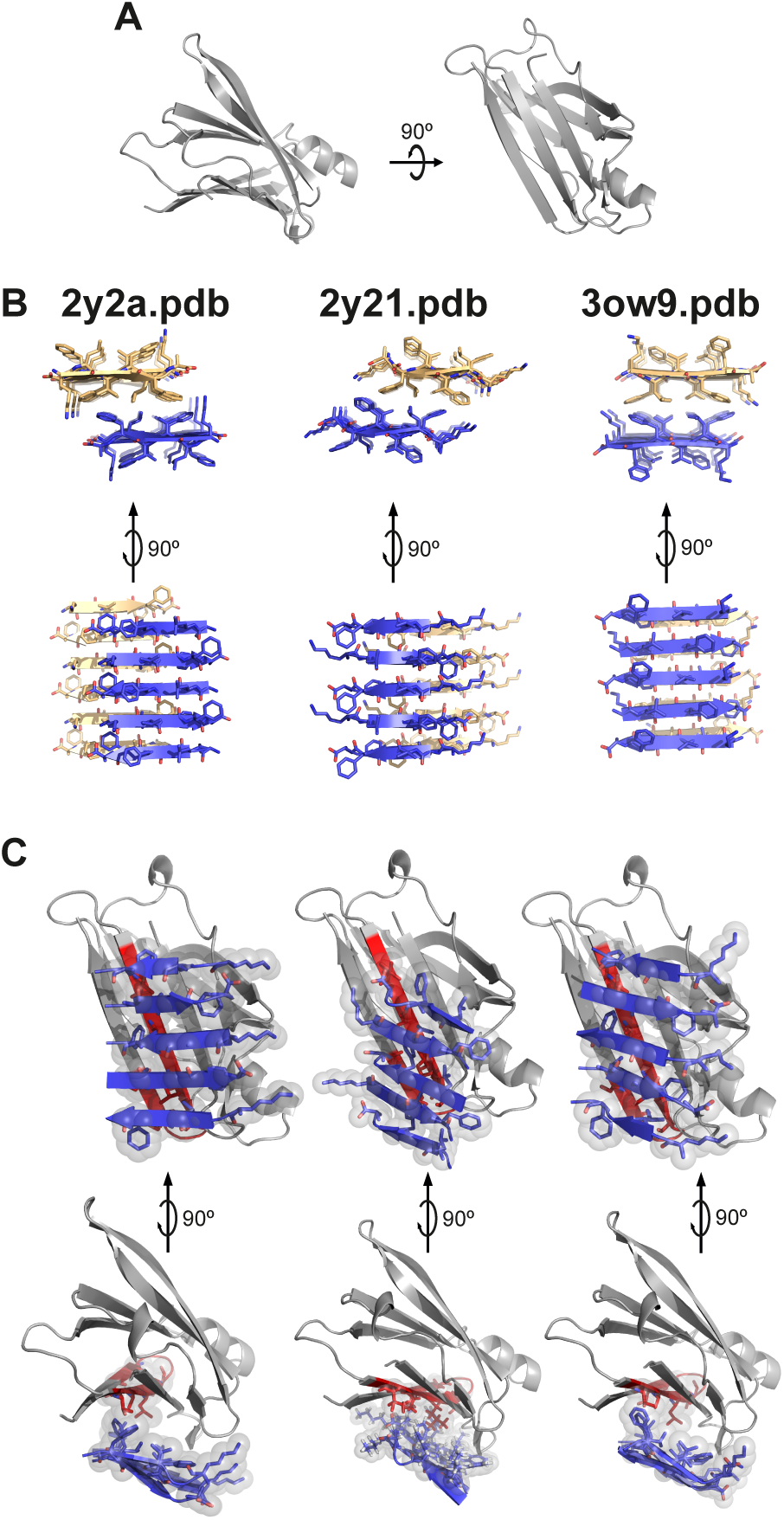
Protein-protein docking of TTR monomer with three KLVFFA polymorphs. Protein-protein docking was performed using monomeric TTR and three KLVFFA polymorphs identified previously (17): 2y2a.pdb, 2y21.pdb and 3ow9.pdb. **(A)** Monomeric TTR obtained from chain A of the model 4TLT.pdb is shown in grey as secondary structure. On the left, view down the hydrophobic pocket. On the right, lateral view. **(B)** Fibrillar structures of KLVFFA from 2y2a.pdb (left), 2y21.pdb (middle), and 3ow9.pdb (right). Only one sheet of each KLVFFA polymer was included in the docking, shown in blue. The resulting docking models, shown in **(C)**, suggest that TTR may interact with Aβ42 through residues 105-117. Monomeric TTR is shown in grey with the segment TTR(105-117) in red. Top raw, lateral view of interface. Bottom raw, view down the binding interface. Residues involved in the interaction between monomeric TTR and KLVFFA are shown as sticks. Spheres represent the van der Waals radii of the side chain atoms of the tightly packed binding interface.

The peptide TTR(105-117) was found to be highly amyloidogenic. We first analyzed the amyloidogenicity of TTR sequence by ZipperDB, which measures the propensity of every six-residue segment to form amyloid fibrils (19, 20) (Supplementary Fig.2A). TTR predictions show that the region that contains this segment is highly prone to form amyloid structures. In fact, we found the segment TTR(105-117) and the shorter peptide TTR(106-117) to be amyloidogenic in solution (Supplementary Fig. 2B), which may explain the lack of inhibitory effect in previous studies (21). In contrast to the amorphous aggregates generated by recombinant full-length TTR, these peptides form amyloid-like fibrils (Supplementary Fig. 2B). We then explored several sequence modifications to increase solubility, by eliminating the first tyrosine or adding a charged tag to the N-terminal end. The sequences and names of all the analyzed peptides are listed in Table 2. We found that TTR(105-117) amyloidogenicity was fully hindered by the addition of a poly-arginine tag (Supplemental Fig. 2B).

**Table 2.**
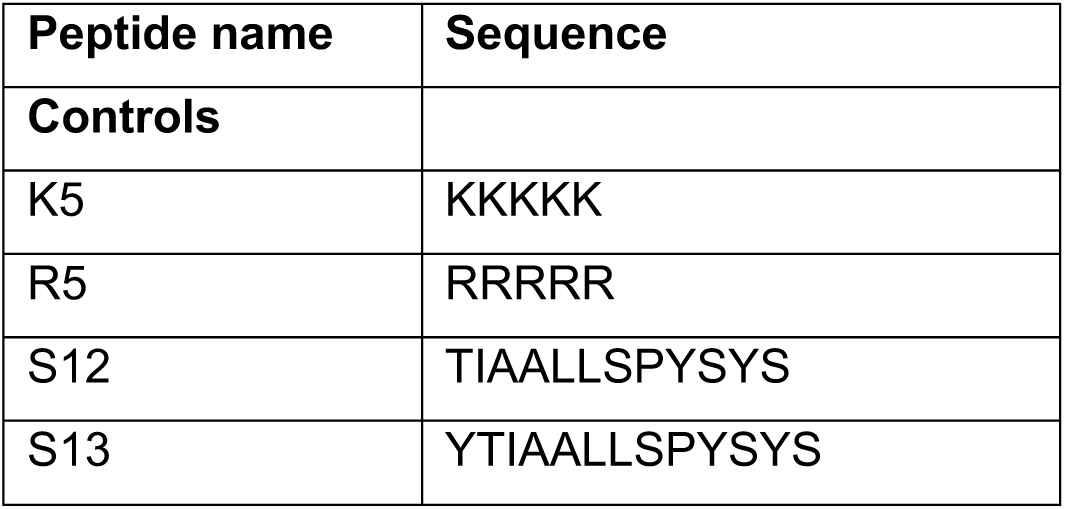
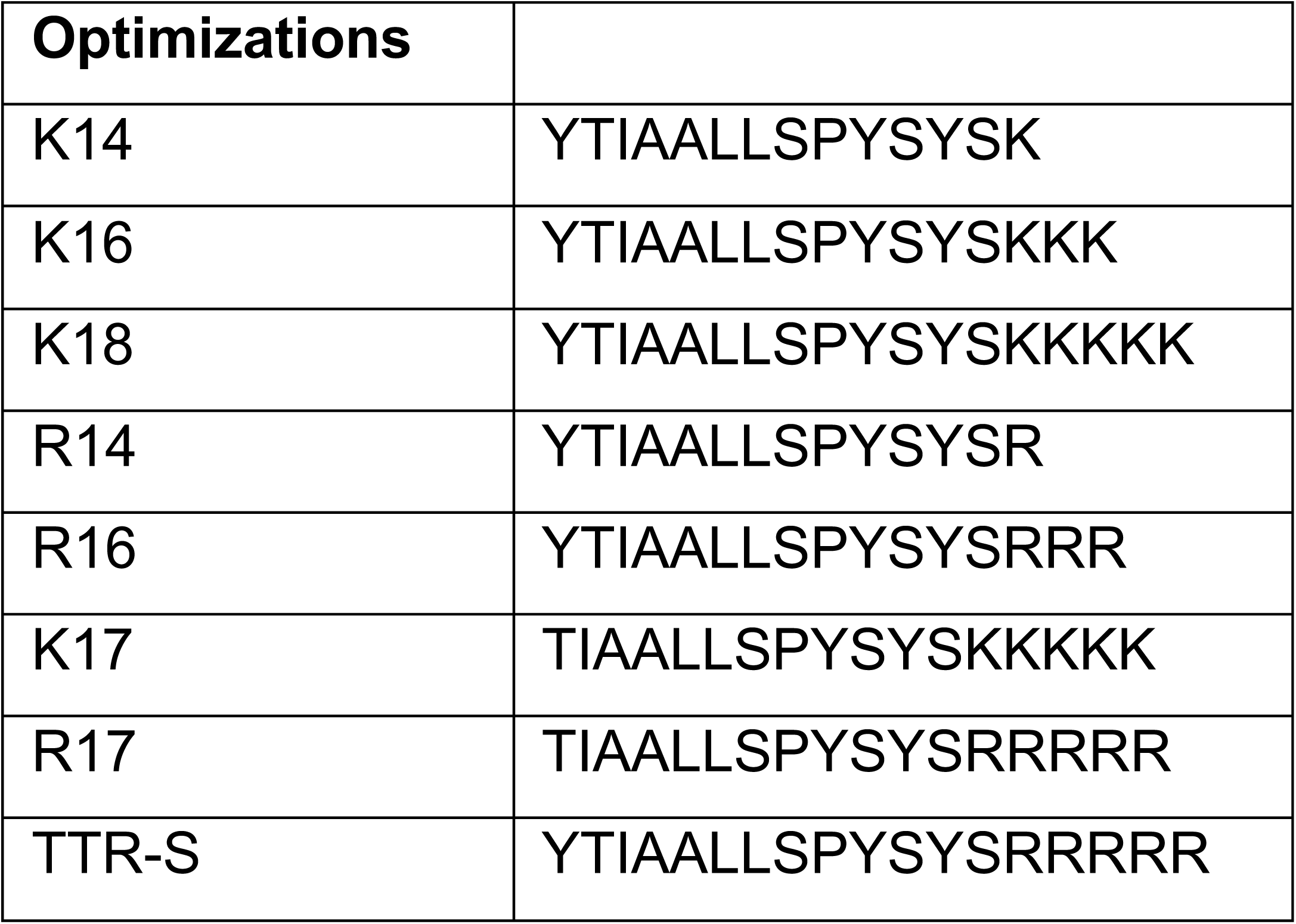
List of peptides evaluated in this study.

| **Peptide name** | **Sequence** |
| --- | --- |
| **Controls** |  |
| K5 | KKKKK |
| R5 | RRRRR |
| S12 | TIAALLSPYSYS |
| S13 | YTIAALLSPYSYS |

| Optimizations |  |
| --- | --- |
| K14 | YTIAALLSPYSYSK |
| K16 | YTIAALLSPYSYSKKK |
| K18 | YTIAALLSPYSYSKKKKK |
| R14 | YTIAALLSPYSYSR |
| R16 | YTIAALLSPYSYSRRR |
| K17 | TIAALLSPYSYSKKKKK |
| R17 | TIAALLSPYSYSRRRRR |
| TTR-S | YTIAALLSPYSYSRRRRR |

The soluble tagged derivatives of the segment TTR(105-117) inhibited Aβ42 fibril formation (Fig. 3 and Supplementary Fig. 3). We first evaluated the inhibitory effect of TTR-derived peptides over Aβ42 fibril formation in thioflavin T (ThT) assays (Fig. 3A and Supplementary Fig. 3). Out of all peptides that we analyzed, the best Aβ42 inhibitory effect was obtained with TTR-S (sequence YTIAALLSPYSYSRRRRR), which contains a 4-arginine tag followed by TTR(105-117) (Supplementary Fig. 3). After an incubation of 2 days, we analyzed the samples by electron microscopy, and found that the addition of TTR-S promoted the formation of amorphous aggregates (Fig. 3B) that were not birefringent when stained with Congo-red (Fig. 3C). Remarkably, we found that TTR-S also promoted the formation of amorphous species when incubated with preformed Aβ42 fibrils (Fig. 3B). These aggregates were also found to be thioflavin T negative (Fig. 3A), and non-birefringent when stained with Congo-red (Fig. 3C). The structural characterization of Aβ42 amorphous aggregates by circular dichroism showed that the addition of TTR-S to preform fibrils resulted in a significant structural shift from beta to helical secondary motifs (Fig. 3D). Additionally, we found that this structural modification results in cytotoxicity protection (Fig. 4). HeLa, PC12, and SH-5YSY cells were subjected to Aβ42 in the absence and presence of TTR-S at different molar ratios, and cell metabolic activity was followed by MTT reduction (Fig. 4A-D). We found that the addition of TTR-S results in a significant reduction of Aβ42 cellular toxicity. This protective effect may be explained by the inhibition of the formation of A11-positive oligomers as a result of the incubation with TTR-S (Fig. 4D). It is worth noting that the larger aggregates did not resolve in the SDS-page gel.

**Figure 3.**
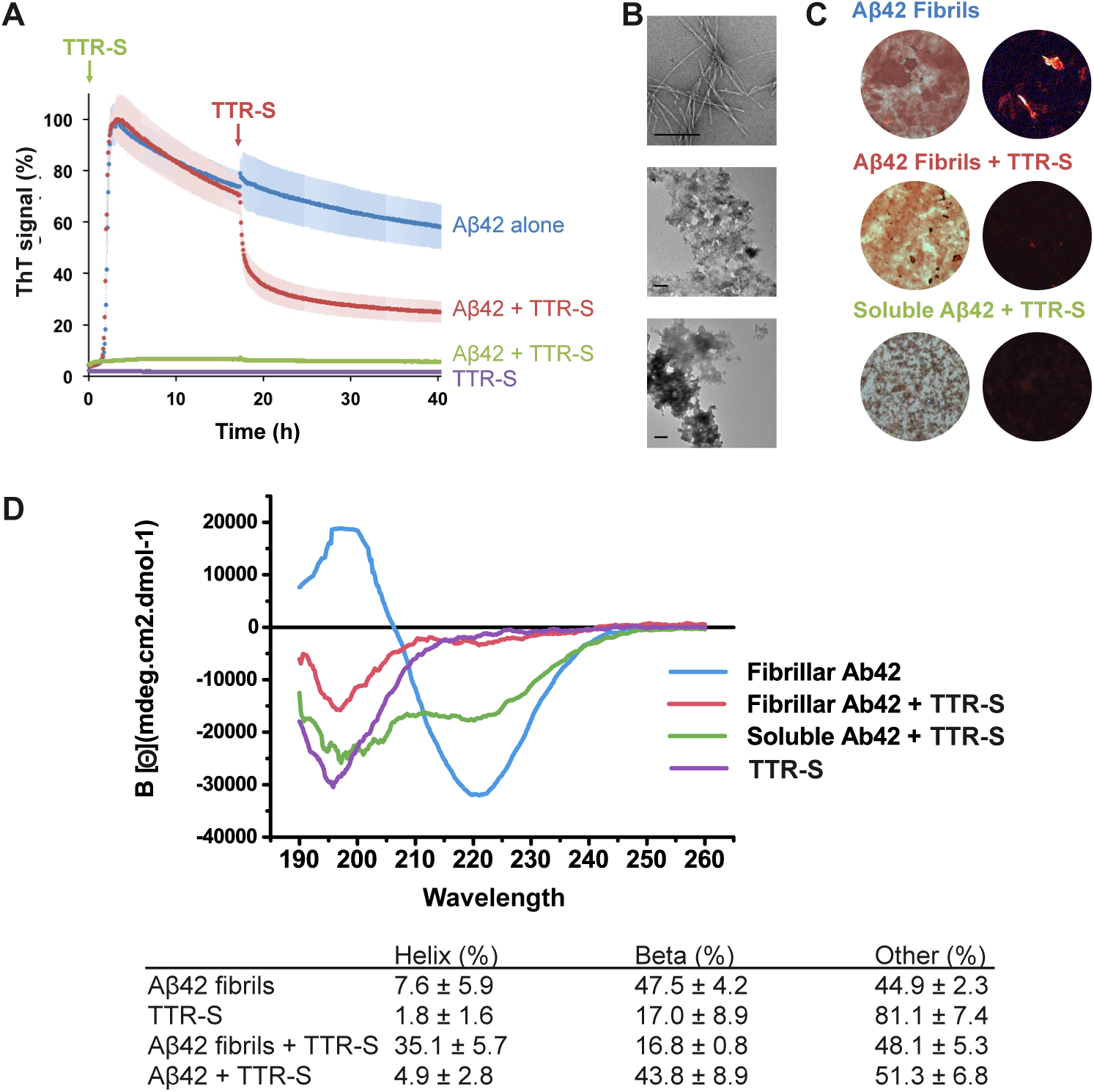
Effect of TTR-S on both soluble and fibrillar Aβ42. **(A)** Thioflavin T (ThT) fluorescence was measured from samples containing 10 μM Aβ42 in the presence and absence of a 3-fold molar excess of TTR-S. TTR-S was added before incubation (green), or after 18 hours of incubation (red). TTR-S alone (purple) and buffer (not shown) were used as negative controls. n=3. Error bars, S.D. For a complete assessment of Aβ42 inhibition by TTR-derived peptides, see Supplementary Fig. 4. **(B)** Electron micrographs of Aβ42 fibrils formed in the absence of inhibitor (up), Aβ42 amorphous aggregates formed when TTR-S was added after 18 hours of incubation (middle), and Aβ42 amorphous aggregates formed when TTR-S was added prior to incubation (bottom). Scale bar, 400 nm. **(C)** Congo red staining of samples shown in ***A*** under bright field (left) and polarized light (right). **(D)** Circular dichroism traces of Aβ42 fibrils, Aβ42 fibrils after addition of TTR-S, and soluble Aβ42 after addition of TTR-S. Below, the table shows the calculated percentage of various structural conformations: helix, beta-stranded, and others (turns and unstructured). These analyses indicate that the TTR-derived peptide TTR-S inhibits Aβ42 aggregation and promotes the formation of non-amyloid amorphous aggregates.

**Figure 4.**
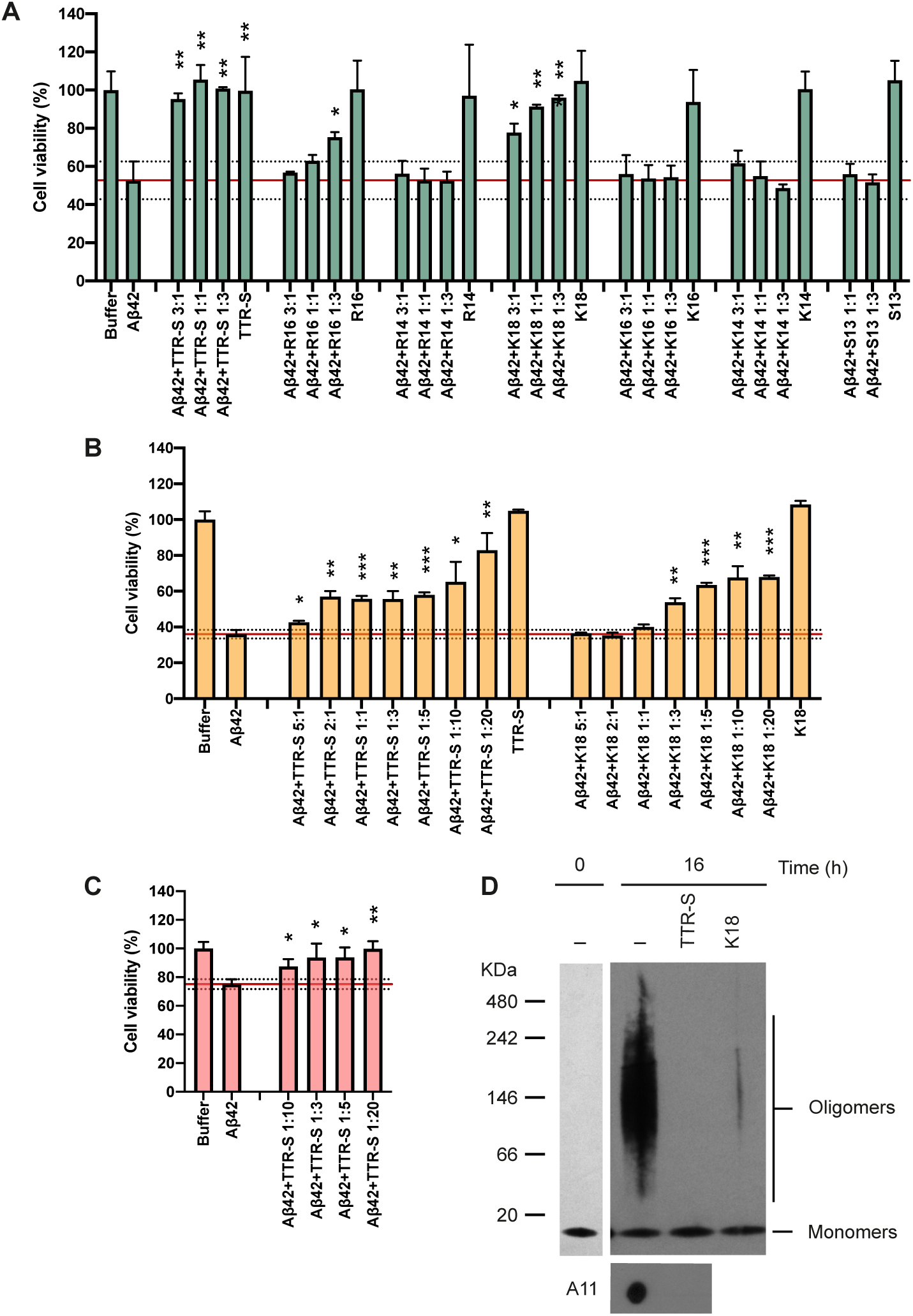
TTR-S inhibition of Aβ42 cytotoxicity. **(A-D)** Cell viability assay followed by MTT reduction in where HeLa **(A)**, PC12 **(B)**, and SH-5YSY **(C)** cells were treated with 10 μM Aβ42 in the absence and the presence of increasing concentrations of peptides. Molar ratios of Aβ42 and peptides are labeled. As a negative control, cytotoxicity of peptides alone was also measured. Buffer-treated cells were considered 100% viability and used for normalization. Cell toxicity measured in the absence of peptides is marked with a red continuous line (mean) and dashed lines (S.D.) n=3. Error bars, S.D. *p ≤ 0.05, **p ≤ 0.005, ***p ≤ 0.0005, N.D., significance not detected. **(D)** Non-denaturing electrophoresis followed by immunoblot of Aβ42 after 16 hours of incubation at 37 °C without and with TTR-S. The same samples were analyzed by dot blot with the oligomer-specific A11 antibody (bottom). These results suggest that the inhibition of Aβ42 cytotoxicity by TTR-S results from the elimination of toxic oligomeric species.

Next we found that the binding of TTR-S to soluble and fibrillar Aβ42 alters their protease resistance. Others have shown that TTR increases Aβ42 clearance *in vivo* (1). We reasoned that the large assemblies that form upon binding to TTR might be more sensitive to proteolytic activity than amyloid fibrils, thereby facilitating clearance. This hypothesis would also explain the protective effect found *in vivo* (6). We explored this hypothesis by analyzing proteolytic sensitivity of TTR-S-derived amorphous aggregates (Fig. 5). We incubated soluble and fibrillar Aβ42 with TTR-S over night, and collected the insoluble fractions by centrifugation. The immuno-dot blot of insoluble fractions showed that amorphous aggregates that result from the incubation of TTR-S with both soluble and fibrillar Aβ42 were easily digested by proteinase K after one hour (Fig. 5A and 5B). In contrast, Aβ42 fibrils showed a significantly higher resistance to proteinase K digestion. We included soluble Aβ42 in the assay as a control of proteolytic activity.

**Figure 5.**
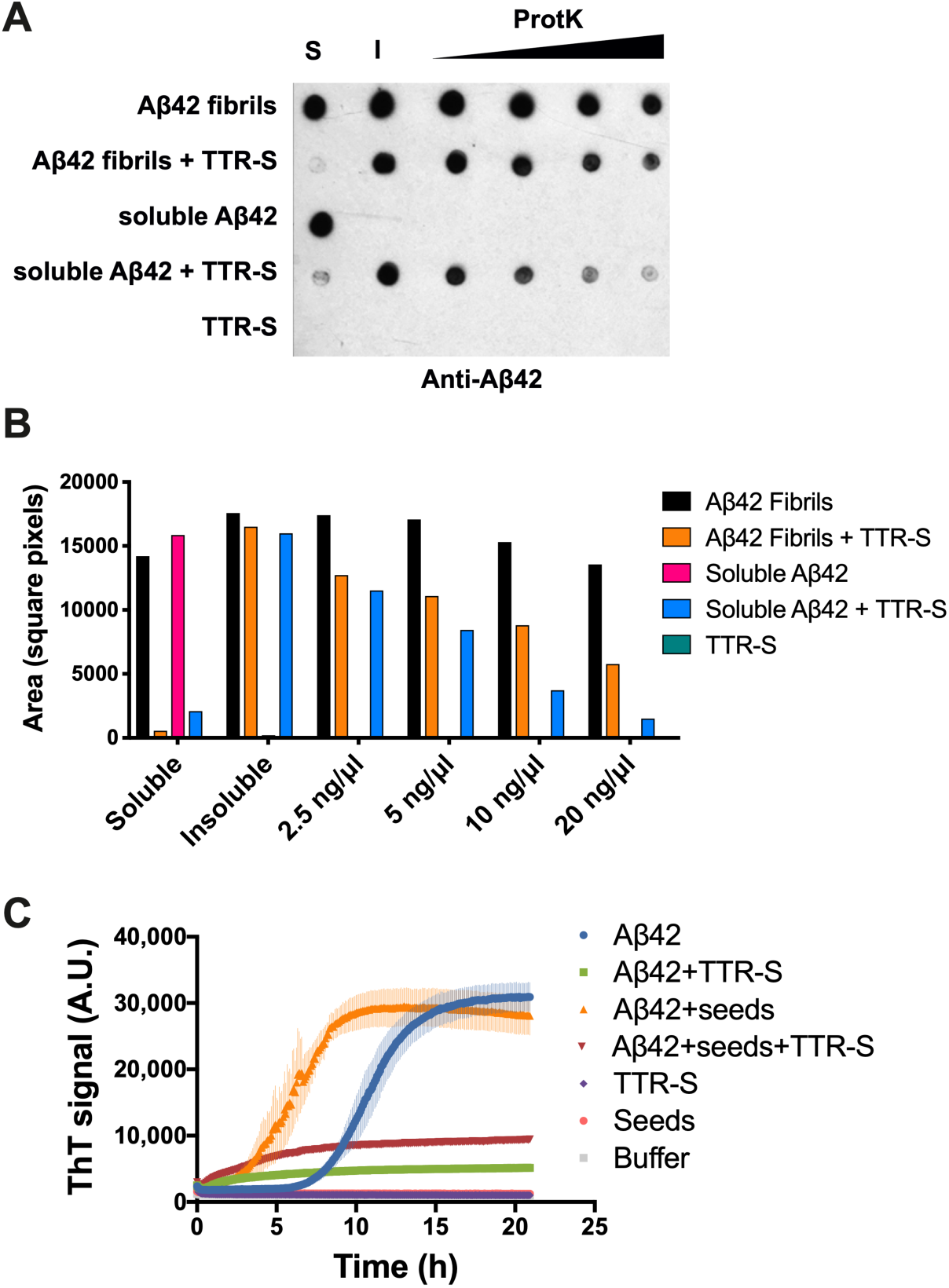
Evaluation of the inhibitory effect of TTR-S by protease digestion and amyloid seeding. **(A)**Anti-Aβ42 immuno-dot blot of Aβ42 aggregates after proteinase K digestion. Soluble Aβ42 and pre-formed fibrils were incubated with TTR-S. Half of the sample was subjected to centrifugation and soluble (S) and insoluble (I) fractions were collected. Increasing concentrations of proteinase K were added to the other half of the sample and incubated at 37° C for one hour. Samples were analyzed by dot blot using E610 specific anti-Aβ42 antibody. As negative control, we included a sample with TTR-S alone. The signal was quantified using ImageJ and plotted in **(B).** This assay shows that Aβ42 amorphous aggregates generated after the addition of TTR-S are degraded by proteinase K more readily than Aβ42 fibrils. **(C)** Amyloid seeding assay followed by ThT signal. Soluble Aβ42 was incubated with AD *ex-vivo* seeds in the presence or absence of TTR-S. Samples with only TTR-S, only AD seeds or buffer were included as negative control. Note that the addition of TTR-S results in inhibition of Aβ42 seeding. n=3. Error bars, S.D.

Finally, we evaluated TTR-S ability to hinder amyloid seeding caused by fibrils extracted from the brain of an AD patient (Fig. 5C). The extraction of Aβ fibrils was performed by several cycles of homogenization and ultracentrifugation as described by Tycko and colleagues (22). Amyloid seeding was measured by thioflavin T fluorescence. As expected, we found that the addition of sonicated patient-derived fibrils to soluble Aβ42 resulted in the acceleration of fibril formation. In contrast, incubation with TTR-S resulted in full inhibition of Aβ42 fibril formation even in the presence of patient-derived Aβ seeds.

## Discussion

Our results indicate that the dissociation of the TTR tetramer is required to prevent cytotoxicity from Aβ oligomers. These results are consistent with the model proposed by Buxbaum and coworkers in where dissociated TTR monomers efficiently bind Aβ oligomers, which are thought to be responsible for cellular Aβ-associated toxicity (16, 23). They also found that tetrameric TTR binds to soluble monomeric Aβ, thereby suppressing aggregation *in vitro* (16). Others have also found an association between genetic stabilization of transthyretin and decreased risk of cerebrovascular disease (24). Consistently, an additional study found that the stabilization of the tetrameric form of TTR promotes Aβ clearance in a mouse model of Alzheimer’s disease (25). In our study, we did not find any effect of tetrameric TTR on Aβ42 cytotoxicity. For instance, the tetrameric non-amyloidogenic T119M variant did not inhibit Aβ42 cytotoxicity (Fig. 1). Consistently, the non-amyloidogenic variant L55P/T119M did not inhibit cytotoxicity whereas the amyloidogenic L55P variant did (Supplementary Fig. 1). Taken together, other studies and our results suggest that tetrameric TTR may be protective *in vivo* perhaps not by recruiting toxic oligomers, but by sequestering and promoting clearance of Aβ monomers. In contrast, dissociated TTR seems to inhibit Aβ cytotoxicity by binding to toxic oligomeric species triggering the formation of large non-toxic assemblies.

Using computational tools, we generated a TTR-derived peptide that inhibits Aβ42 fibril formation and cellular toxicity (Fig. 3). This TTR-derived peptide, here called TTR-S, contains a poly-arginine tag followed by the highly amyloidogenic segment TTR(105-117) (Supplementary Fig. 3). Our results are consistent with previous NMR studies that show that the interaction of TTR residues Lys15, Leu17, Ile107, Ala108, Ala109, Leu110, Ala120, Val121 to Aβ contributes to the inhibition of Aβ fibril formation *in vitro* (16). In addition, others have shown that the segment TTR(106-110), one residue shorter, was capable of binding Aβ when immobilized on a membrane (8). However, the same peptide was not effective inhibiting Aβ aggregation in solution (21), unless the peptide was made cyclic therefore not amyloidogenic (26), probably because of the high amyloidogeneicity of this segment (Supplemental Fig. 3). In our study, we solubilized TTR(105-117) by adding a charged tag, which confers higher solubility and also hinders self-aggregation (Supplementary Fig. 3). The addition of a poly-arginine tag was sufficient to convert a highly amyloidogenic peptide into an inhibitor of Aβ42 cellular toxicity even at substoichiometric concentrations (Fig. 4A).

TTR-S inhibits Aβ42 oligomer formation whilst promoting the formation of non-toxic, non-amyloid, amorphous aggregates. Previous studies of the inhibition of Aβ cytotoxicity by TTR have revealed that TTR co-aggregates with Aβ oligomers into large non-toxic assemblies, thereby inhibiting cellular toxicity *in vitro* (9). TTR-S mimics this effect and promotes the formation of large aggregates (Fig. 3B) that do not bind thioflavin T (Fig. 3A), do not show birefringence upon staining with Congo red (Fig. 3C), and display a non-beta secondary structure when subjected to CD (Fig. 3D). These amorphous species share some resemblance with the large unstructured aggregates found after the addition of EGCG to intrinsically disordered Aβ and other amyloid proteins (27). Similar to EGCG-derived Aβ aggregates, TTR-S-derived aggregates display a unique secondary structure that results in reduced thioflavin T fluorescence and lack of birefringence upon binding to Congo red (Fig. 3A and 3C).

Aβ42 amorphous aggregates generated upon binding to TTR-S are more sensitive to protease digestion (Figs. 4 and 5). As discussed above, TTR inhibits Aβ oligomer toxicity by co-aggregation into large non-toxic assemblies (9). However, those studies did not assess protease sensitivity of Aβ species. We observe that TTR-S-derived aggregates, both from soluble and fibrillar Aβ, are more sensitive to proteinase K than preformed Aβ fibrils. We speculate that TTR may promote Aβ clearance *in vivo* by a similar mechanism; this is, the formation of large assemblies that are more prone to digestion and clearance.

Finally, we evaluated the inhibitory effect of TTR-S on Aβ amyloid seeding. As shown previously, we found that the addition of *ex-vivo* fibril extracts to soluble Aβ accelerates fibril formation (28, 29). In contrast, we found that TTR-S inhibits amyloid seeding catalyzed by Aβ extracted from the brain tissue of an AD patient (Fig. 5C).

In summary, our results suggest that the dissociation of the TTR tetramer into monomers is required to prevent cytotoxicity from Aβ oligomers. In addition, we found that a segment derived from TTR, TTR-S, exerts anti-amyloid activity, thereby inhibiting Aβ42 cytotoxicity and amyloid seeding. We also found that TTR-S binds to soluble and fibrillar species causing a structural rearrangement that leads to an increase of protease sensitivity. Finally, we found that TTR-S inhibits amyloid seeding catalyzed by Aβ fibrils extracted from the brain of an AD patient. The present study represents an expansion of the current knowledge on the mechanism of protection of TTR over Aβ42 cellular toxicity and opens a potential therapeutic avenue.

## Acknowledgments

We thank David Eisenberg for his significant support. We also thank Dr. Harry V. Vinters for supplying the AD brain tissue and the patient who generously donated it. This work was supported by NIA, National Institutes of Health (NIH), Grant RF1 AG048120 (to David Eisenberg), the Amyloidosis Foundation Grants 20160759 and 20170827 (to L.S.); and the People Programme (Marie Curie Actions) of the European Union’s Seventh Framework Programme (FP7/2007–2013) under Research Executive Agency (REA) Grant Agreement 298559 (to L.S.) L.S. is a consultant for ADRx, Inc.

## Author contributions

Qin Cao: Investigation, Validation, and Data Curation – Daniel H. Anderson, Wilson Liang, and Joshua Chou: Investigation – Lorena Saelices: Conceptualization, Methodology, Validation, Formal analysis, Investigation, Data Curation, Writing Original Draft, Writing Review & Editing, Visualization, Supervision, Project administration, and Funding acquisition

## Experimental Procedures

### Patients and Tissue Material

Post-mortem brain tissue from the occipital lobe of an 83-year-old female patient of Alzheimer’s disease was obtained from Dr. Vinters at UCLA Pathology Department. The patient was previously evaluated for the presence of amyloid plaques by immunohistochemistry and pathologically diagnosed for Alzheimer’s disease. The University of California, Los Angeles Office of the Human Research Protection Program granted exemption from Internal Review Board review because the specimen was anonymized.

### Recombinant protein purification

TTR mutants were cloned and purified as previously described (18). Briefly, exponentially growing *E. coli* Rosetta™(DE3)pLysS Competent Cells (Millipore) were treated with 1 mM of IPTG for 3 h. Mutant TTR were purified by Ni-affinity chromatography using HisTrap columns (GE Healthcare), followed by gel filtration chromatography using a HiLoad 16/60 Superdex 75 column (GE Healthcare) running on an AKTA FPLC system. Amyloid-β peptide (Aβ42) was overexpressed through *Escherichia coli* recombinant expression system and was purified as reported previously (30). The fusion construct for Aβ42 expression contains an N-terminal His tag, followed by 19 repeats of Asn-Ala-Asn-Pro, TEV protease site and the human Aβ42 sequence. Briefly, the fusion construct was expressed into inclusion bodies in *E. coli* BL21(DE3) cells. 8 M urea was used to solubilize the inclusion bodies. Fusion proteins were purified through HisTrap HP Columns, followed by Reversed-phase high-performance liquid chromatography (RP-HPLC). After TEV cleavage, Aβ42 peptide was purified from the cleavage solution by RP–HPLC followed by lyophilization. To disrupt preformed aggregation, lyophilized Aβ42 was resuspended in 100% Hexafluoroisopropanol (HFIP) which was finally removed by evaporation.

### Size-exclusion chromatography with multi-angle light scatter and refractometer detection

Components of 0.5 µg protein samples were separated by a silica-based size-exclusion column (Toso Biosep G3000SWXL; 5 µm beads, 7.8×300 mm with guard column). The elution buffer contained 0.025M NaH_2_PO_4_, 0.1 M Na_2_SO_4_, 1mM NaN_3_, weighed on an analytical scale, titrated together to pH 6.5 with NaOH, diluted to final volume in a 2 liter volumetric flask. The flow rate was 0.4 ml/min. Samples were 0.1 µm filtered (Millipore UltraFree MC centrifugal filters). Each injection was 40-50 µl. The peaks were detected by UV absorbance at 280 nm (Waters 2487), light scatter and index of refraction (Wyatt Technologies MiniDAWN and OPTILAB DSP). Molecular weights were calculated with Astra software, from the light scatter and refractometer signals, transferring the dn/dc and normalization parameters determined from the monomer peak of bovine serum albumin.

### TTR aggregation assay

TTR aggregation assays were performed as previously described (31). Briefly, 1 mg/ml TTR sample in 10 mM Sodium Acetate pH 4.3, 100 mM KCl, 10 mM EDTA was incubated at 37° C during a maximum of 7 days. TEM micrographs were taken after 7 days of incubation. Insoluble fractions were sampled and used to follow TTR aggregation by immuno-dot blot.

### Transmission electron microscopy (TEM)

TEM was performed to visualize TTR mutant aggregation and the fibrillation of Aβ42 in presence of TTR mutants or TTR-derived inhibitors. 5 μl solution was spotted onto freshly glow-discharged carbon-coated electron microscopy grids (Ted Pella, Redding, CA). Grids were rinsed three times with 5 μl distilled water after 3 min incubation, followed by staining with 2% uranyl acetate for 2 min. A T12 Quick CryoEM electron microscope at an accelerating voltage of 120 kV was used to examine the specimens. Images were recorded digitally by Gatan 2kX2k CCD camera.

### Cell lines

HeLa, PC-12 Adh, and SH-SY5Y (ATCC; cat. # CRL-2, CRL-1721.1 and CRL-2266, respectively) cell lines were used for measuring the toxicity of Aβ42. HeLa cells were cultured in DMEM medium with 10% fetal bovine serum, PC-12 cells were cultured in ATCC-formulated F-12K medium (ATCC; cat. # 30-2004) with 2.5% fetal bovine serum and 15% horse serum.

### MTT-based cell assay

We performed MTT-based cell viability assay to assess the cytotoxicity of Aβ42 with or without the addition of TTR mutants or TTR-derived peptide inhibitors. A CellTiter 96 aqueous non-radioactive cell proliferation assay kit (MTT) (Promega cat. #G4100, Madison, WI) was used. Prior to toxicity test, HeLa, PC-12, and SH-5YSY cells were plated at 10,000, 15,000, and 10,000 cells per well, respectively, in 96-well plates (Costar cat. # 3596, Washington, DC). Cells were cultured in 96-well plates for 20 hr at 37°C in 5% CO2. For Aβ42 and inhibitors samples preparation, purified Aβ42 was dissolved in PBS at the final concentration of 10 μM, followed by the addition of TTR mutants or TTR-derived inhibitors at indicated concentrations. The mixtures were filtered with a 0.2 μm filter and further incubated for 16 hours at 37°C without shaking for fiber formation. To start the MTT assay, 10 μl of pre-incubated mixture was added to each well containing 90 μl medium. After 24 hours of incubation at 37°C in 5% CO2, 15 μl Dye solution (Promega cat. #G4102) was added into each well. After incubation for 4 hours at 37°C, 100 μl solubilization Solution/Stop Mix (Promega cat. #G4101) was added to each well. After 12 hours of incubation at room temperature, the absorbance was measured at 570 nm with background absorbance recorded at 700 nm. Three replicates were measured for each of the samples. The MTT cell viability assay measured the percentage of survival cell upon the treatment of the mixture of Aβ42 and inhibitors. The cell viability (%) after treatment with Aβ42 with and without TTR-derived peptide inhibitors was calculated by normalizing the cell survival rate using the PBS buffer-treated cells as 100% viability and 2% SDS treated cells as 0% viability.

### Computational Docking of fibrillar KLVFFA and monomeric TTR

Protein-protein docking between monomeric TTR and Aβ42 fibrillar segments was performed using the ClusPro server (32). Monomeric TTR was generated by removing chain B of the asymmetric unit of 4TLT.pdb (18). Three fibrillar polymorphs of the Aβ42 segment KLVFFA were analyzed: 2Y2A.pdb, 2Y21.pdb, and 3OW9.pdb ((17)). To increase binding surface, only one sheet of each polymorph was included in the modeling.

### Thioflavin T fibrillation assay

Purified Aβ42 was dissolved in 10 mM NaOH at the concentration of 300 μM. Aβ42 was diluted into PBS buffer at the final concentration of 30 μM, and was mixed with 30 μM Thioflavin T (ThT) and different concentrations of TTR mutants or TTR-derived peptide inhibitors. The reaction mixture was split into four replicates and placed in a 394-well plate (black with flat optic bottom). The ThT fluorescence signal was measured every 5 min using the Varioskan plate reader (Thermo Scientific, Inc.) or FLUOstar Omega plate reader (BMG Labtech) with excitation and emission wavelengths of 444 and 484 nm, respectively, at 37°C.

### Western and Immuno-Dot Blot

The aggregation of His-tagged TTR mutants was followed by immuno-dot blot analysis as described in Saelices et al. 2015 using SuperSignal^®^ West HisProbe™ Kit following manufacturer’s instructions (Life Technologies). Briefly, 100 μl of samples was spun at 13,000 rpm during 30 minutes, the pellet resuspended in the same volume of fresh buffer, and spun again. The final pellet was resuspended in 6M guanidine chloride and dotted onto nitrocellulose membranes (0.2 μm, Bio-Rad). We used a concentration of the HisProbe antibody of 1:10,000. The aggregation of Aβ42 was followed by western and immuno-dot blot analysis. For the western blots, 7.5 μl of 6 μM samples were separated by SDS-PAGE (NuPAGE 4-12% Bis-Tris Gel, Life Technologies) or native gels (NativePAGE 4-16% Bis-Tris Gel, Life Technologies) and transferred onto a nitrocellulose membrane (iBlot^®^ 2 NC Mini Stacks, Life Technologies) by iBlot^®^ 2 membrane transfer system (Life Technologies). For the Immuno-dot blot analysis, 2 to 15 µl of samples were dotted onto a nitrocellulose membrane (Bio-RAD). We used a concentration of 6E10 antibody of 1:2000 (Biolegend®, (33)), OC antibody of 1:25,000 and A11 antibody of 1:500 (Millipore and Life Technologies, respectively; (34)). The HRP conjugated anti-mouse IgG antibody (Sigma-Aldrich) was used as secondary antibody at a concentration of 1:2000, and anti-rabbit IgG antibody (Thermo Scientific) was used to detect OC and A11 at a concentration of 1:1000. The membrane was finally developed using the SuperSignal^®^ West Pico Chemiluminescent Substrate Kit (Life Technologies), following manufacturer’s instructions.

### Circular Dichroism (CD)

Secondary structures of Aβ42 samples were analyzed by CD spectroscopy. Samples (200 µl) were placed into a 1-mm path length quartz cell (Hëllma Analytics). A Jasco J-810 UV-Vis spectropolarimeter was employed. Spectra were obtained in a wavelength range of 190 to 250 nm, with a time response of 2 s, a scan speed of 50 nm/min, and a step resolution of 0.1 nm. Each spectrum was the average of five accumulations. All the samples were assayed at a concentration of 50 µg/ml in 15 mM phosphate buffer (pH 7.4). Spectra were recorded at 25° C. The results are expressed as the mean residue molar ellipticity,

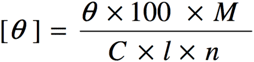

Where *θ* is the ellipticity in degrees, *l* is the optical path in cm, *C* is the concentration in mg/ml, *M* is the molecular mass and *n* is in the number of residues in of the sample molecule. The ellipticity of the samples of Aβ after addition of TTR-S was normalized by using the ellipticity of TTR-S as a blank. The percentage of the various structural conformations was calculated by using the software package CDPro, which includes three programs to determine the secondary structure fractions: CONTIN, SELCON, and CDSSTR (35). The results are displayed as the mean value and standard deviation of the three methods.

### Proteinase K assay

Soluble Aβ42 was prepared in PBS and filtered (0.22 nm). Aβ42 fibrils were obtained after 1-2 weeks of incubation in PBS at RT. Three-fold molar excess of TTR-S was added to soluble or fibrillar Aβ42 and incubated for 16 hours. Soluble and insoluble fractions were extracted by centrifugation of half of the sample prior to incubation. PK digestion was performed in a reaction volume of 100 μl containing 10 μM of Aβ42 samples, and varying concentrations of PK (from 0.2 μg/ml to 50 μg/ml) in 0.1 M Tris-HCl buffer, pH 7.5 for 1 h at 37 °C. Reactions were quenched with 1 mM PMSF, and samples were analyzed dot blot using E610 specific anti-Aβ42 antibody. As negative control, we included a sample with TTR-S alone. The signal was quantified using ImageJ.

### Statistical Analysis

Statistical analysis of cell assays was performed with Prism 7 for Mac (GraphPad Software) using an unpaired t test. All samples were included in the analysis. All quantitative experiments are presented as independent replicates or as means ± SD of at least three independent experiments.

---

**Figure S1.**
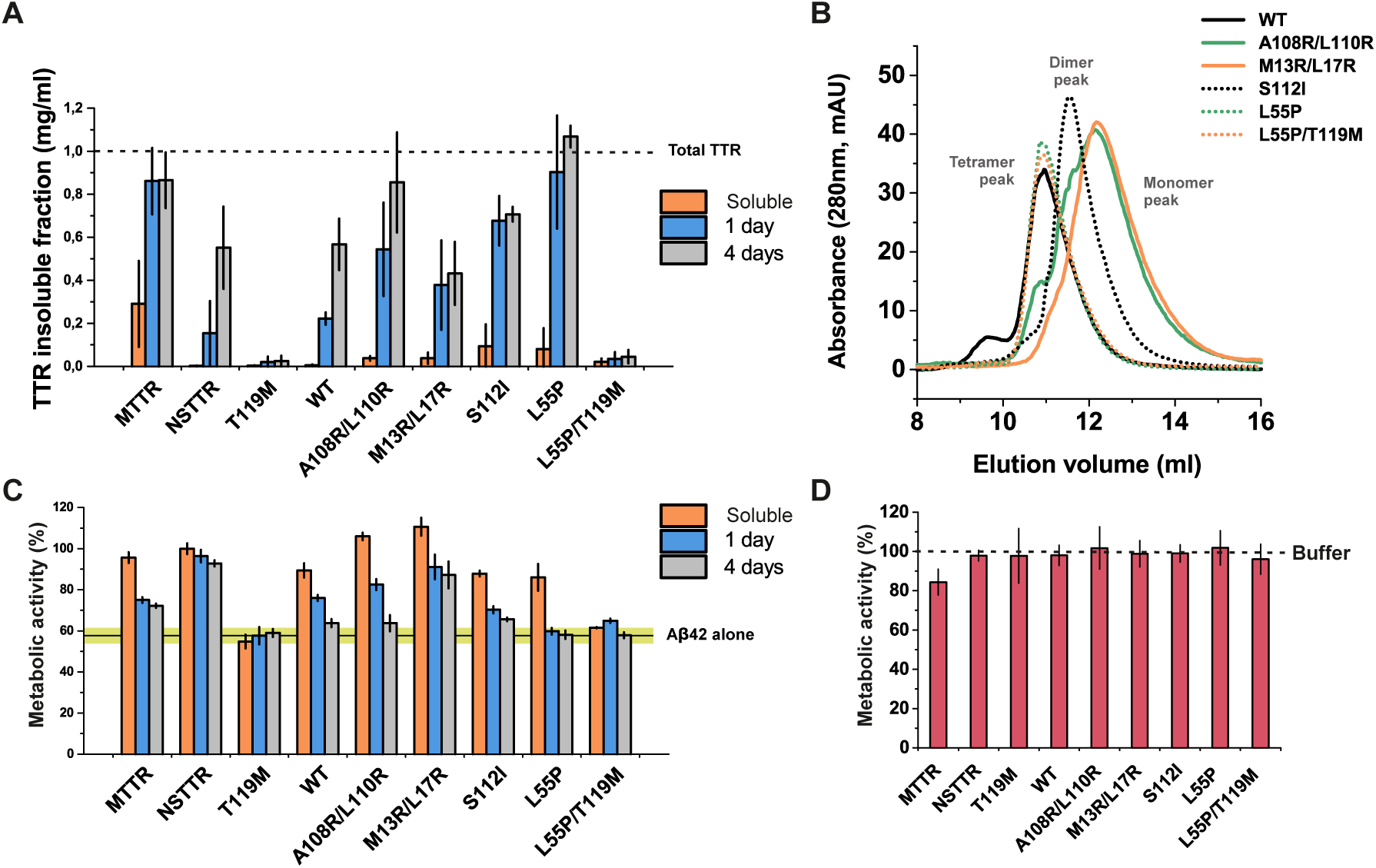
Inhibition of Aβ42 cytotoxicity by TTR variants. **(A)** Aggregation assay of TTR variants followed by protein concentration of insoluble fractions collected prior incubation, or after 1, and 4 days of incubation at 37° C, as labeled. **(B)** Size exclusion chromatography of soluble TTR variants at pH 7.4. **(C)** Cytotoxicity assay of Aβ42 in the presence of TTR variants at different stages of aggregation, followed by MTT reduction. A five-fold molar excess of soluble TTR, and aggregated samples collected after 1 and 4 days of incubation was added to soluble 10 μM Aβ42 and incubated overnight. Samples were added to HeLa cells, and MTT reduction was measured after 24 hours. Buffer-treated cells were considered 100% viability and used for normalization. **(D)** Cytotoxicity of TTR variants in the absence of Aβ42 measured by MTT reduction. TTR final concentration was kept the same as for **(C)**, 50 μM.

**Figure S2.**
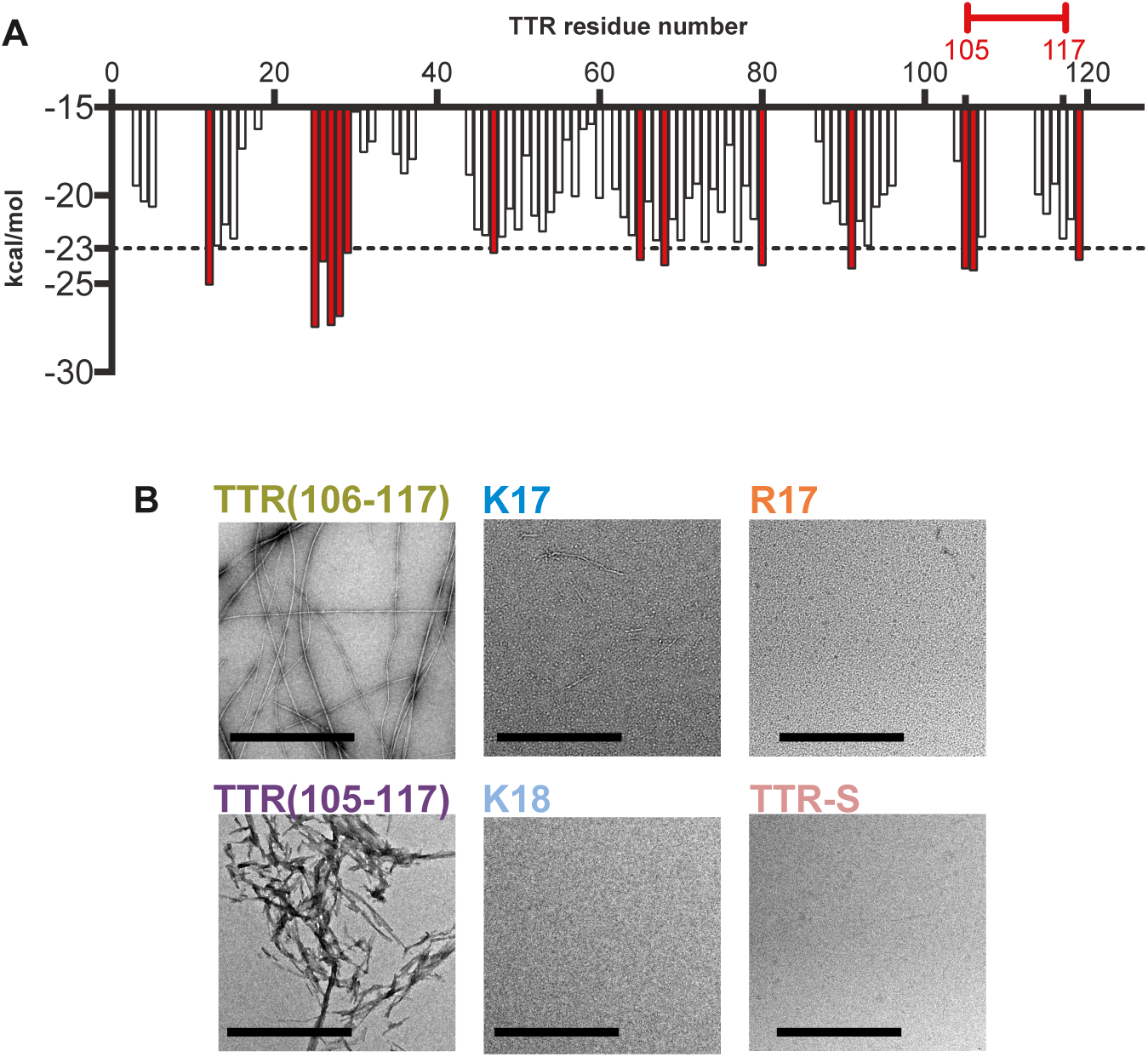
Evaluation of TTR-derived peptides in the absence of Aβ42. **(A)** Propensities of steric zipper formation of each 6-residue segment within the TTR sequence. Computationally predicted segments with steric zipper propensities are represented with *red bars*. The segment identified from docking modeling is marked with a red bracket. **(B)** Electron micrographs of TTR-derived peptides after 26 hours of incubation at 37°C. Scale bar, 200 nm.

**Figure S3.**
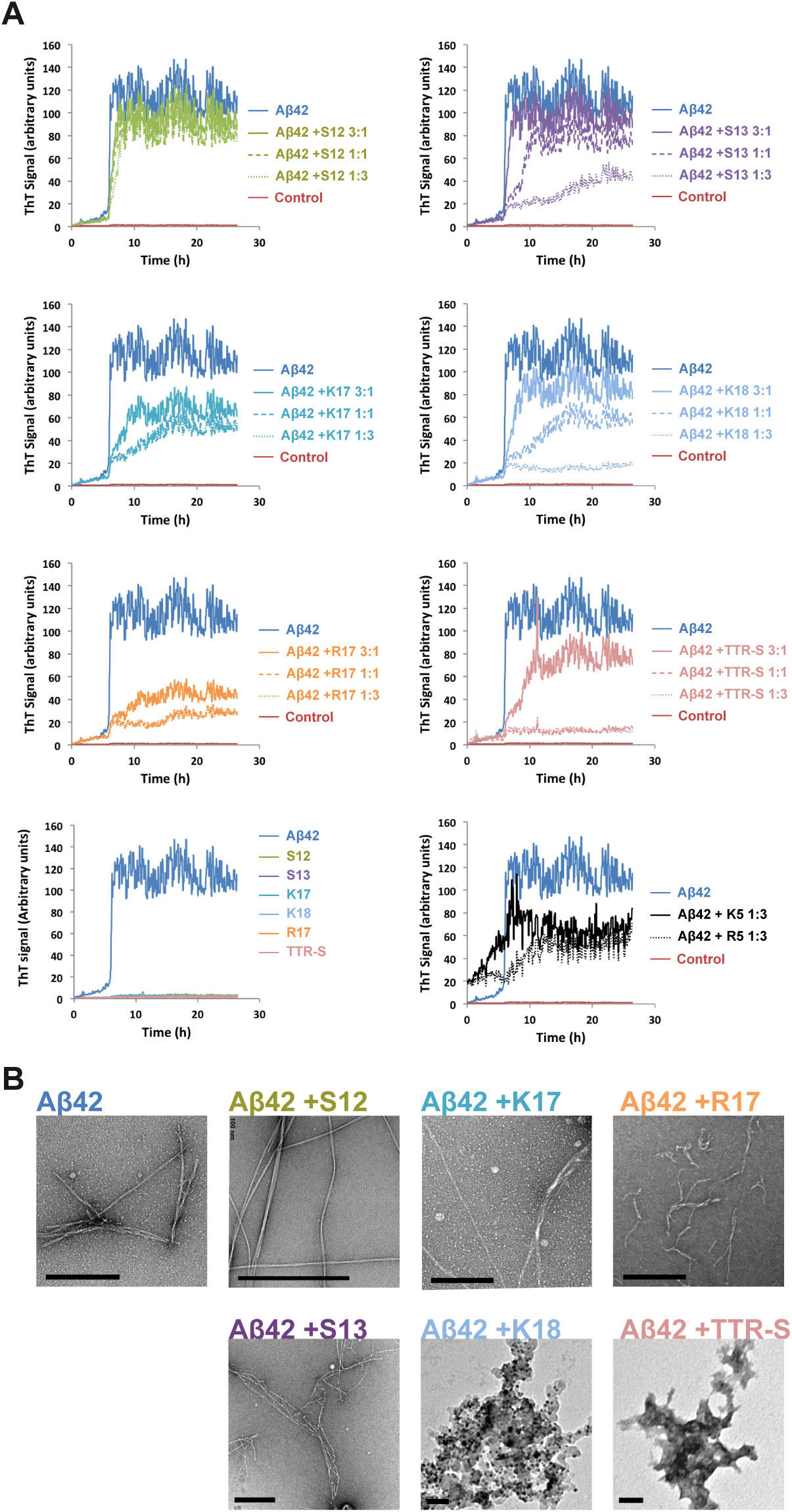
Evaluation of the inhibitory effect of TTR-derived peptides. **(A)** Thioflavin T (ThT) fluorescence was measured from samples containing 10 μM Aβ42 in the presence and absence of TTR-derived peptides at several molar ratios, as labeled. TTR-derived peptides were added at the starting point. Peptide sequences are listed in Table 2. Buffer was used as negative control. **(B)** Electron micrographs of samples collected from ***A***, incubated for 26 hours in the presence and absence of 3-fold molar excess of peptides, as labeled.

**Figure S4.**
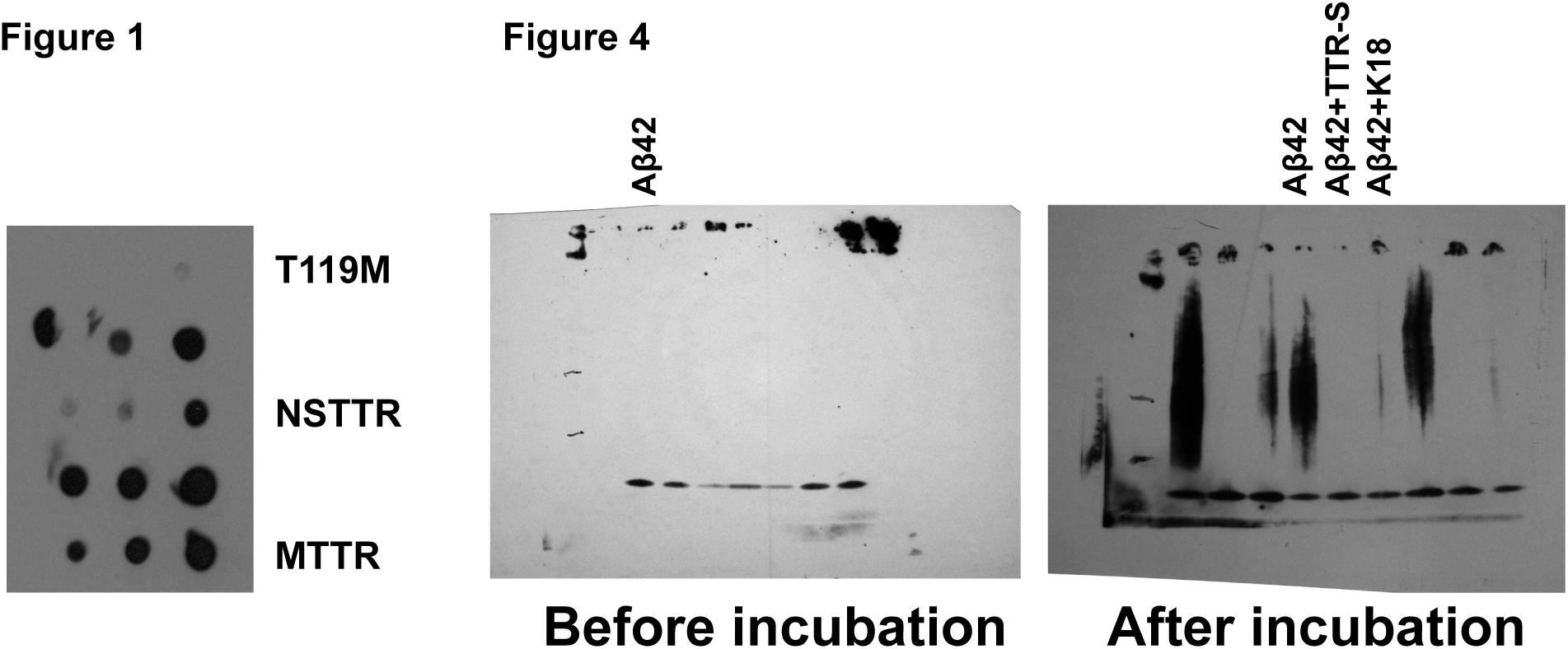
Uncut Gels.

